# Phase Contrast Tomography (PCT)

**DOI:** 10.1101/2024.03.17.585445

**Authors:** Ying Ma, Wenjing Feng, Yunze Lei, Lin Ma, Juanjuan Zheng, Sha An, Min Liu, Peng Gao

## Abstract

Phase contrast microscopy has been employed for 2D imaging of thin samples since its invention. Herein, we propose and demonstrate phase contrast tomography (PCT) that incorporates the scanning of LED illumination with quantitative phase contrast microscopy (QPCM) to realize 3D phase imaging of a sample. The proposed PCT is demonstrated 3D imaging of 200-nm polystyrene microspheres (PMs) and sub-organelles inside COS7 cells. The results reveal that the proposed PCT has high spatial resolution and stability without speckle noise, and thence, it has a great potential to be applied to industrial testing and life science research.

## 1. Introduction

Phase contrast imaging, which can convert the phase distribution of a transparent object into intensity modulation, is widely used in studies of biological structures [1-3]. The basic principle of phase contrast is to introduce a certain phase-shift on the unscattered component of an object wave with respect to the scattered component. Due to the common-path configuration, the phase contrast method has the advantage of being insensitive to environmental vibration. Popescu et al. proposed spatial light interference microscopy (SLIM) [4], which uses annular illumination and a spatial light modulator (SLM) to retarder the phase of the unscattered light on a ring on the Fourier plane. Notably, the phase distribution recovered by these techniques is two dimensional, i.e., the average of the 3D refractive index (RI) distribution of a sample along the axial direction. As nearly all the samples have 3D-distributed internal structures, 3D phase imaging is pivotal to accurately detect sample’s structures without affecting their states.

Optical diffraction tomography (ODT) is the pioneer of the 3D phase imaging techniques to obtain 3D refractive index distribution of a sample. ODT incorporates scanning illumination into an off-axis DHM to acquire the 2D complex amplitude transmittance of the sample along different directions, from which the 3D RI of the sample can be obtained. ODT has been rapidly developed and widely applied in life science research [5-7]. Thereinto, Cotte’s [8] and Chen’s groups [9] further improved the spatial resolution of the traditional ODT to visualize the sub-organelles inside live cells. However, the mechanical scanning in the traditional ODT invites mechanical vibration and limits the imaging speed to some extent. Accordingly, the structured illumination generated by a digital micromirror device (DMD) or a spatial light modulator (SLM) is coupled into the traditional ODT to ensure the temporal resolution [10-12]. And the ODT with low coherence noise was realized by using a temporally low-coherence source coupled into two DMDs [13] or an SLM [14]. Furthermore, Chowdhury et al. improved the stability of the ODT by combining the structured illumination generated by an SLM with the broadband common-path digital holographic microscopy [15,16]. Notably, the polarized diffraction phase imaging technique [17] has been proposed to adjust and balance the energy ratio of reference wave to object wave, optimizing the fringe contrast of the hologram as well as the reconstruction accuracy.

In aside of ODT, researchers have developed several advanced techniques to achieve high-quality 3D label-free imaging with high stability and low noise, for instance, using low-coherence shear interference [18-20], partially coherent illumination engineering [21], diffraction-based approaches with multiple LED illuminations [22-25], and nearly in-line holography with an LED source [26]. These methods are straightforward and cost-effective. Yet, currently, there is still lack of an imaging technique that can image subcellular organelles, such as mitochondria, in 3D and with high spatial resolution. Although the white-light diffraction tomography based on a SLIM system has been proposed, it still needs to record a z-stack of raw data that covers the entire sample and relies on a complex deconvolution process to obtain the 3D distribution of the sample [27]. Meanwhile, the high-quality imaging of subcellular organelles remains unrealized [27].

This study innovatively proposes phase contrast tomography (PCT). PCT combines the scanning illumination scheme of ODT with quantitative phase contrast microscopy, realizing the high-resolution, high-stability, and low-noise 3D label-free imaging of a sample. The proposed PCT was demonstrated with 3D imaging of 200-nm-diameter polystyrene microspheres (PMs) and sub-organelles inside COS7 cells, including mitochondria and lipid droplets.

## 2. Results

### 2.1 Construction of PCT

The schematic diagram of PCT is shown in **Fig. 1(a)**. 25 identical LEDs evenly distributed on a ring were turned on one by one to illuminate the sample at different angles in a time sequence. The illumination vector of each illumination beam was determined in advance by installing a temporary lens (Appendix 5.1). Under each illumination angle, the sample is imaged by a telescope system comprised of a microscopic objective (OBJ) and a tube lens (TL). The spectra of the generated object wave appear in the middle Fourier plane of OBJ-TL, which is comprised of unscattered and scattering terms (high- and low-frequency components). A spatial light modulator (SLM) was placed on this plane to retard the phase of the unscattered term and low-frequency term of the object wave (red area in **Fig. 1(a)**) with phase values of 0, 0.5π, π, and 1.5π. As the illumination annularly scans, the phase-shifted phase contrast images were recorded by a sCMOS camera. **Fig. 1(b)** shows the data processing workflow of PCT. First, the 2D high-frequency scattering fields of the sample were obtained through a simple phase-shifting reconstruction regime [28]. Second, the 3D frequency spectrum of the sample’s scattering potential was determined by projecting the spectrum of these 2D high-frequency scattering fields to the corresponding 3D Ewald crowns and stitching them together. Eventually, the 3D quasi-RI map of the sample was calculated after performing the Wiener filtering processing and an inverse Fourier transform. With this technique, the 3D distribution of a sample can be revealed despite not the real 3D RI distribution, which highlights the high-frequency structures of the sample in 3D (removing the phase modulation from the bulky cell body). Throughout the manuscript, the obtained 3D quasi-RI distribution is normalized between 0 and 1. The partially coherent illuminations from 25 LEDs endow PCT with high-resolution and noise-free characteristics. Meanwhile, the structure of common-path interference configuration makes the system very immune to external disturbances (Appendix 5.2: illumination stability of the PCT). Thus, the PCT has unprecedent advantages on stability and robustness. In particular, the organelles-specific feature (benefiting from the removal of spatially slow-varied phase from the cell body) of the high-quality 3D structures of sub-organelles inside COS7 cells, including lipid droplets and mitochondria, are rendered, as shown in **Fig. 1(b)**.

**Fig. 1.**
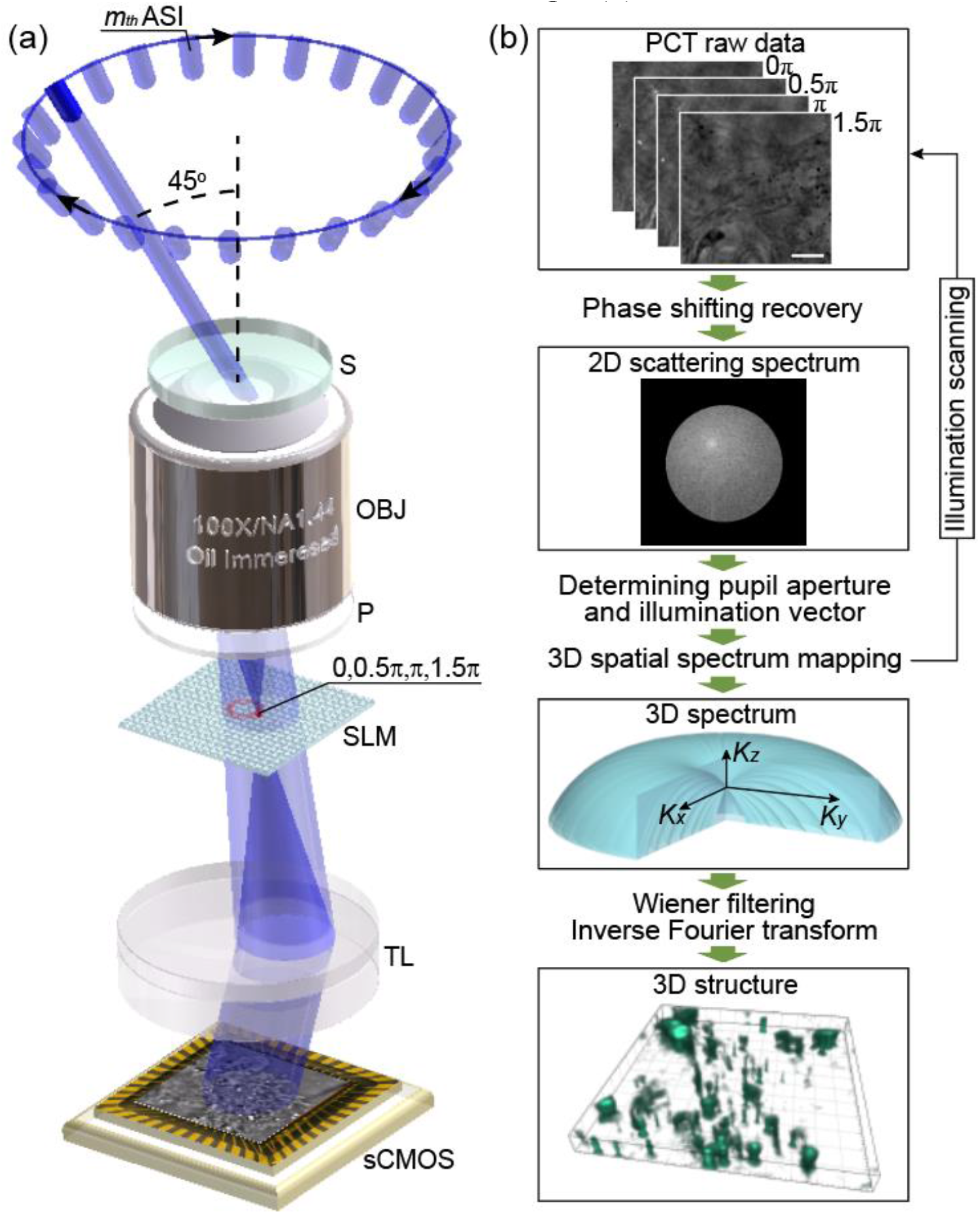
Imaging principle of the proposed PCT. (a) The schematic diagram of PCT. (b) The workflow of 3D reconstruction of PCT. ASI, annular scanning illumination; OBJ, objective lens; P, polarizer; SLM, spatial light modulator; sCMOS, scientific complementary metal-oxide-semiconductor; S, sample; TL, tube lens. The scale bar in (b) represents 5 μm.

### 2.2 The 3D imaging of 200-nm polystyrene microspheres (PMs)

To verify the 3D imaging capability and to identify the spatial resolution of the proposed PCT, single-layer dense 200-nm PMs (RF200C, Huge Biotechnology, China) were prepared and imaged by the PCT system. The sample was sequentially illuminated with oblique illumination featuring a polar angle of ∼ 45° and an azimuth angle of 2mπ/25, with *m*=0, 2, …, 24. And, four phase-shifted intensity images were recorded for each illumination. The 3D image of the 200-nm PMs was reconstructed by the proposed PCT, as shown in **Fig. 2. Figure 2(a)** shows the x-y section of the sample at z = 0 μm plane. **Figure 2(b)** illustrates the enlarged view of a 200-nm PM marked with a white box in **Fig. 2(a)**. Moreover, **Fig. 2(c)** demonstrates the y-z section of the 200-nm PM along the white dotted line in **Fig. 2(a)**. Due to the limited spectrum coverage along the axial direction, the 3D structure distribution of the 200-nm PM has an axially elongated profile. **Figure 2(d)** depicts the line profiles along the lateral and axial directions through the center of a 200-nm PM. Further, the spatial resolution was determined by measuring the full width at half maximum (FWHM) of the line profiles of 30 individual PMs. **Figure 2(e)** shows the statistics of the lateral resolution and the axial resolution among the 30 PMs. Consequently, the lateral resolution of the proposed PCT was 270 ± 9 nm while the axial one was 890 ± 42 nm (mean ± s. d.), which was in accordance with the performance (lateral resolution: 230 nm and the axial resolution 860 nm) of low-coherence ODT based on off-axis DHM [14]. Based on the parallel control algorithm (Appendix 5.3), the proposed PCT achieved a volumetric imaging of 40×110×40 μm^3^ at an imaging speed of 0.8 Hz and a remarkable spatial resolution (lateral 270 ± 9 nm and axial 890 ± 42 nm), and the imaging speed can be further improved by reducing the exposure time of the sCMOS camera or the imaging field of view.

**Fig. 2.**
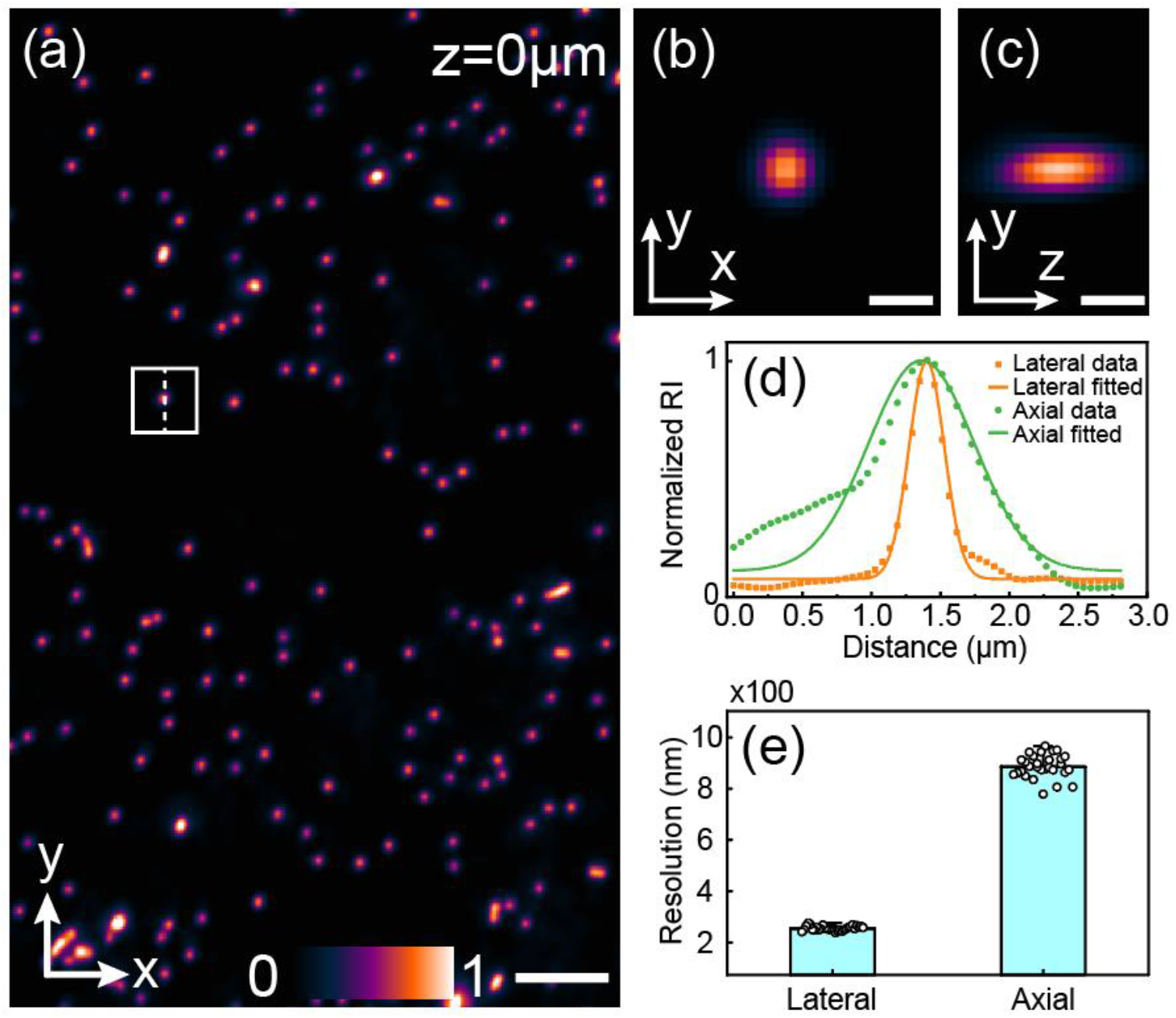
3D imaging performance of the proposed PCT demonstrated by imaging single-layer dense 200-nm PMs. **(a)** The image of the x-y section at z = 0 μm plane. **(b)** Enlarged view of a 200-nm PM marked with a white box in **(a). (c)** The y-z section of a 200-nm PM along the white dotted line in **(a). (d)** Lateral and axial line profiles through the center of a 200-nm PM. **(e)** Statistics of spatial resolution in terms of FWHM among 30 PMs. Scale bars in **(a), (b)**, and **(c)** are 2 μm, 0.3 μm, and 0.3 μm, respectively.

To further verify the feasibility of the 3D imaging of the proposed PCT, 3D sparsely distributed 200-nm PMs were prepared using the agarose gel and was imaged by the proposed PCT without the axial scanning, as shown in **Fig. 3. Figure 3(a)** reveals the structure distribution of 200-nm PMs at z = 0 μm plane, showing in-focus PMs with high-contrast and defocused PMs with low contrast. Here, the uneven background was induced by the curdled agarose gel. The in-focus PMs (indicated by the yellow arrows) appear in the x-y section at z = -2.1 μm plane (**Figs. 3(b))**, rather than in the plane of z= +0.1 μm **(Fig.3(c))**. Similarly, other focused PMs (indicated by the cyan arrows) appear in the x-y section at z = +0.1 μm plane (**Figs. 3(f))**, rather than in the plane of z=-2.1 μm (**Figs. 3(e))**. Further, **Figures 3(d), 3(g)** and **3(h)** represent the y-z sections along the blue dotted line in **Fig. 3(b)**, red dotted line in **Fig. 3(f)**, and white dotted line in **Fig. 3(a)** respectively, further confirming that different 200-nm PMs were in focus on different axial positions through the 3D structure recovery by the proposed PCT.

**Fig. 3.**
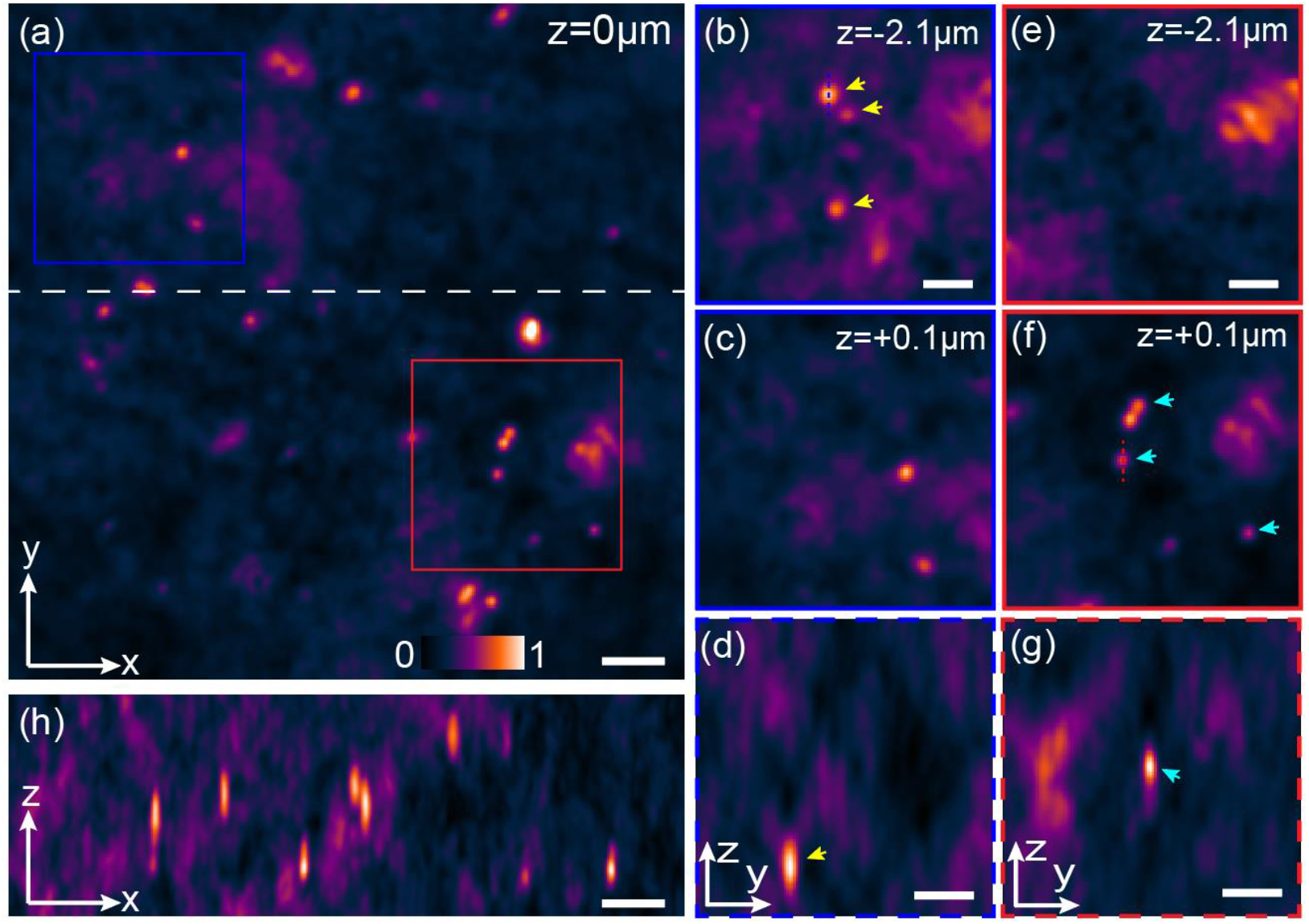
PCT imaging of 3D-distributed 200-nm PMs. **(a)** The x-y section of 200-nm PMs at z = 0 μm plane. **(b–c)** The x-y sections of the blue box in **(a)** on z = -2.1 μm and z = +0.1 μm planes. **(d)** The y-z section along the blue dotted line in **(b). (e–f)** The x-y sections of the red box in **(a)** on z = -2.1 μm and z = +0.1 μm planes. **(g)** The y-z section along the red dotted line in **(f). (h)** The x-z section along the white dotted line in **(a)**. Scale bars in **(a), (b), (d), (e), (g)** and **(h)** are 2 μm, 1 μm, 1.5 μm, 1 μm, 1.5 μm, and 2 μm respectively.

### 2.3 3D imaging of transparent COS7 cells

Furthermore, the proposed PCT was applied to 3D imaging of fixed COS7 cells. **Figure 4(a)** depicts the x-y structure distribution of a fixed COS7 cell on z = -0.3 μm plane, wherein several organelles were identified, including nucleolus inside the nucleus and lipid droplets (LDs). To further manifest the 3D imaging capability of the proposed PCT, we analyzed the structure distribution of the LDs from different perspectives. **Figures 4(b)-(d)** represent the x-y structure distributions of the region marked with a red rectangle in **Fig. 4(a)** at three different z planes. The results show that three different LDs, indicated by a yellow arrow in **Fig. 4(b)**, a cyan arrow in **Fig. 4(c)**, and a green arrow in **Fig. 4(d)**, were in focus at three different z planes, respectively. Moreover, **Fig. 4(e)** demonstrates the x-z section along the red dotted line in **Fig. 4(a)**, where the LDs were asymmetrically stretched along the z-axis due to the limited axial resolution and the aberration of the system. Meanwhile, several LDs distributed on different z positions are shown appropriately in the x-z section, further indicating the 3D imaging capability of the proposed PCT. In addition, we elucidated the 3D structure distribution of two LDs in the region marked with a blue box in **Fig. 4(a)**, as shown in **Figs. 4(f)-(i)**. The LD indicated by a white arrow in **Fig. 4(f)** was in focus on z = 1.9 μm plane, while that indicated with a black arrow in **Fig. 4(h)** was in focus on z = 1.1 μm plane. **Figure 4(i)** is the y-z section along the blue dotted line in **Fig. 4(a)**, showing two LDs located on different axial positions.

**Fig. 4.**
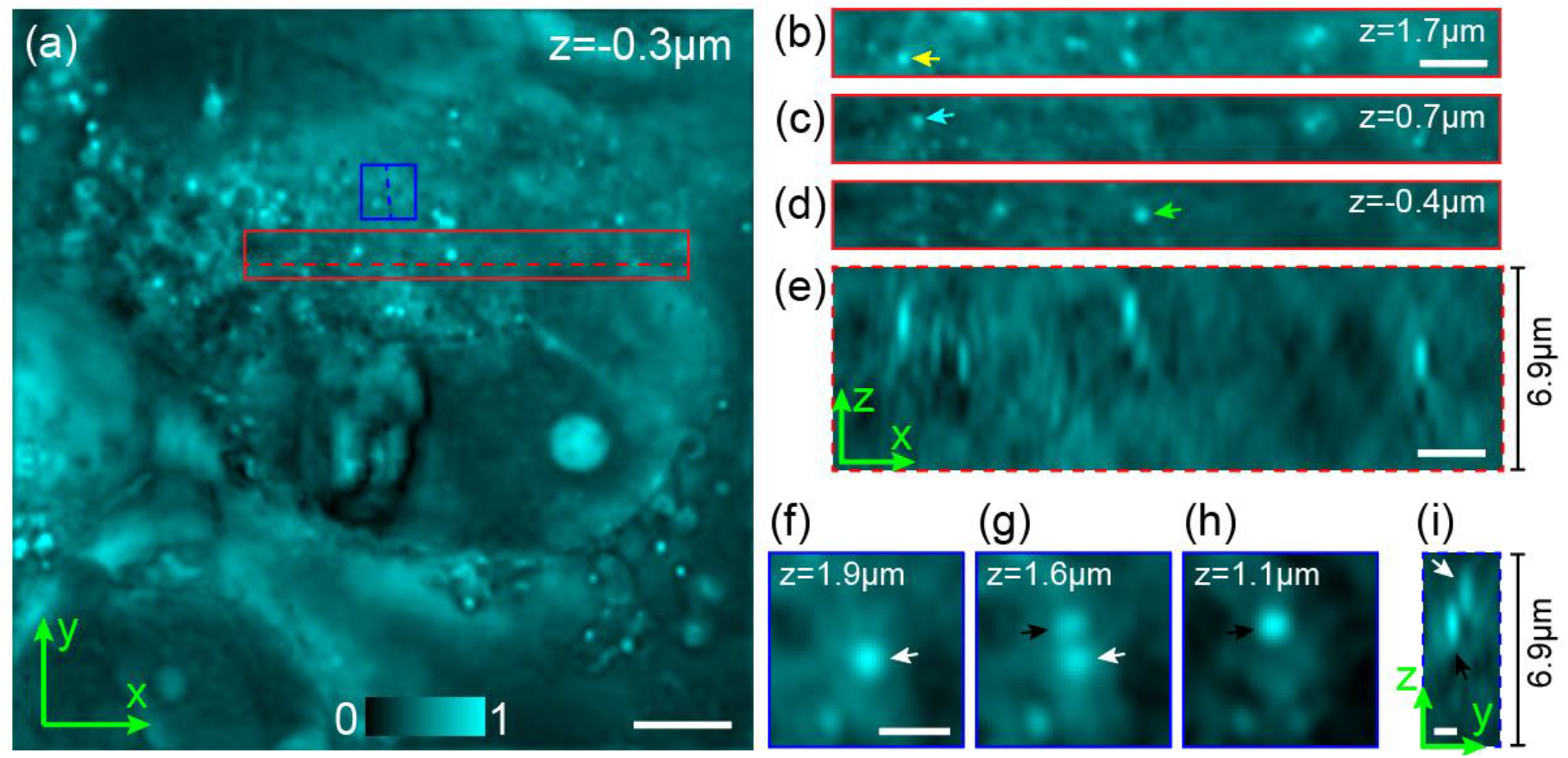
3D label-free imaging of a fixed COS7 cell using PCT. **(a)** The x-y structure distribution at z = -0.3 μm plane. **(b–d)** The x-y structure distributions of the red rectangle in **(a)** on three different z planes. **(e)** The x-z section along the red dotted line in **(a). (f–h)** The x-y structure distributions of the blue box in **(a)** on three different z planes. **(i)** The y-z section along the blue dotted line in **(a)**. Scale bars in **(a), (b), (e), (f)** and **(i)** are 5 μm, 2.4 μm, 2.4 μm, 1 μm, and 0.8 μm, respectively.

Eventually, the proposed PCT was used to visualize simultaneously multiple sub-organelles inside fixed COS7 cells, as shown in **Fig. 5. Figures 5(a)** and **5(b)** demonstrate the traditional bright-field image and the 3D structure distribution of the same COS7 cell in the same field of view (FOV), respectively. Apparently, sub-organelles inside transparent COS7 cells cannot be identified from the traditional bright-field image (**Fig. 5(a)**). Conversely, the 3D structure distribution rendered by the proposed PCT can visualize several sub-organelles, like mitochondria and LDs, inside transparent COS7 cells with high contrast and resolution (**Fig. 5(b)**). The comparison here implies that the proposed PCT indeed allows for the high-resolution, high-stability, and noise-free 3D label-free imaging of sub-organelles inside transparent cells without axial scanning.

**Fig. 5.**
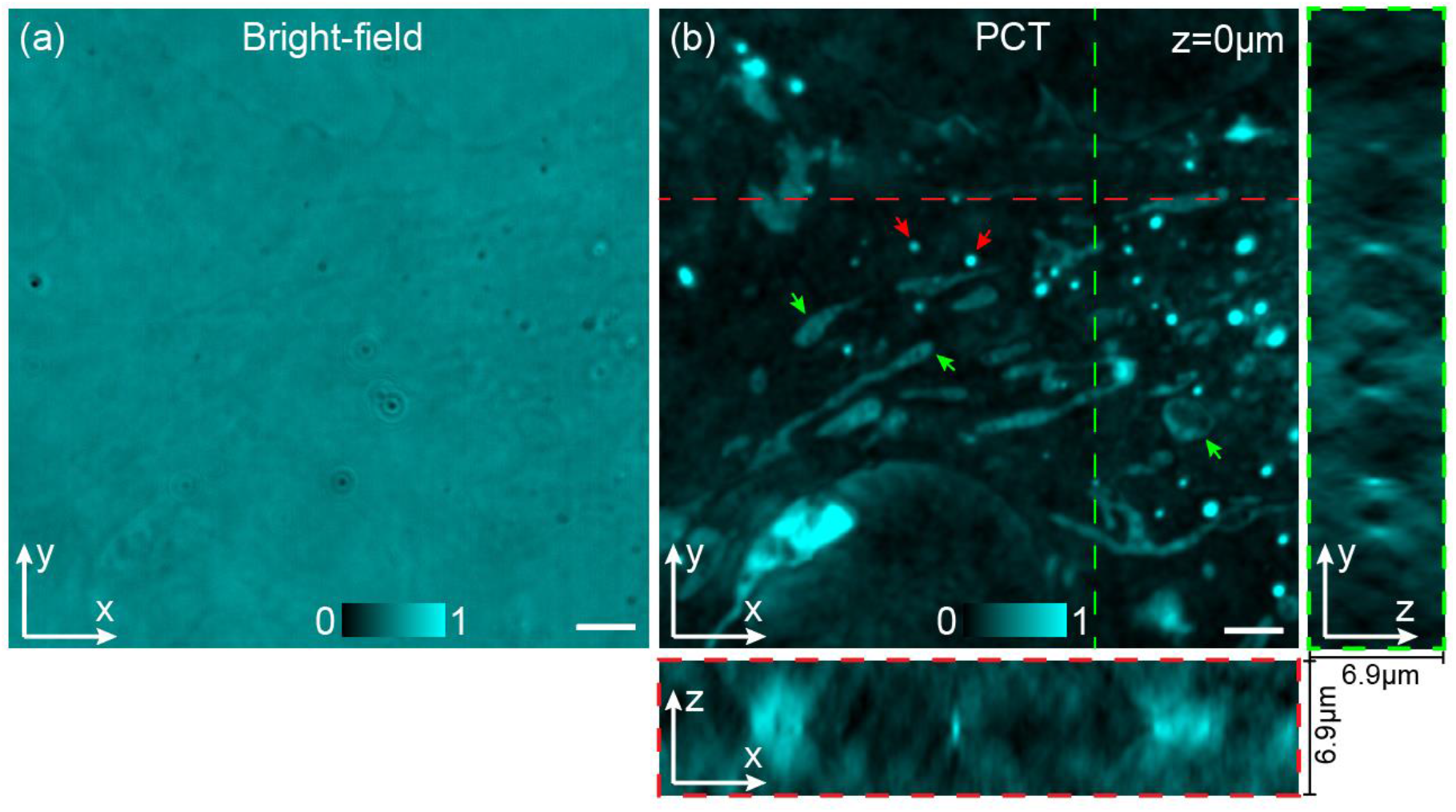
3D imaging of mitochondria and LDs in a fixed COS7 cell using PCT. **(a)** Bright-field image of the fixed COS7 cell, obtained by adding up the 0π intensity image at each illumination angle. **(b)** The x-y, x-z, y-z sections of the fixed COS7 cell in the same field of view (FOV) with **(a)** obtained by using PCT. Organelles pointed by green and red arrows are mitochondria and LDs, respectively. Scale bars in **(a)** and **(b)** are 3 μm.

## 3. Methods

**Figure 6(a)** depicts the schematic diagram of the proposed PCT that was built on a Leica microscope body (DMi8, Leica, Germany). 25 identical LEDs (470 ± 20 nm) were evenly positioned on a ring installed 5 cm above the sample positioned at the focal plane of the objective lens (OBJ; 100×/1.44, Leica, Germany), as shown in **Fig. 6(a)**. As the LEDs were turned on one by one, the sample was sequentially illuminated with partially coherent illuminations in 25 directions, which share a polar angle of 45° and differentiate with an azimuth angle of 2mπ/25 with *m*=0, 2, …, 24. Under each illumination angle, an object wave 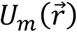 comes into being when the illumination light passes through the weakly scattering sample with a refractive index (RI) distribution of 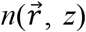. Here, 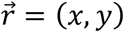 is the lateral coordinate vector. According to the Fourier diffraction theorem [5], 3D real RI distribution of the sample can be obtained by filling the frequency spectrum of 2D scattering fields 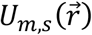 into the 3D frequency spectrum regions (Ewald crowns) of the scattering potential 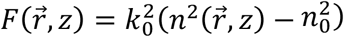 of the sample. Here, *k*_0_ = 2π/λ refers to the wave number in vacuum and λ is the central wavelength of the illumination light. *n*_0_ is the RI of the surrounding medium, and *m* represents the illumination serial number.

**Fig. 6.**
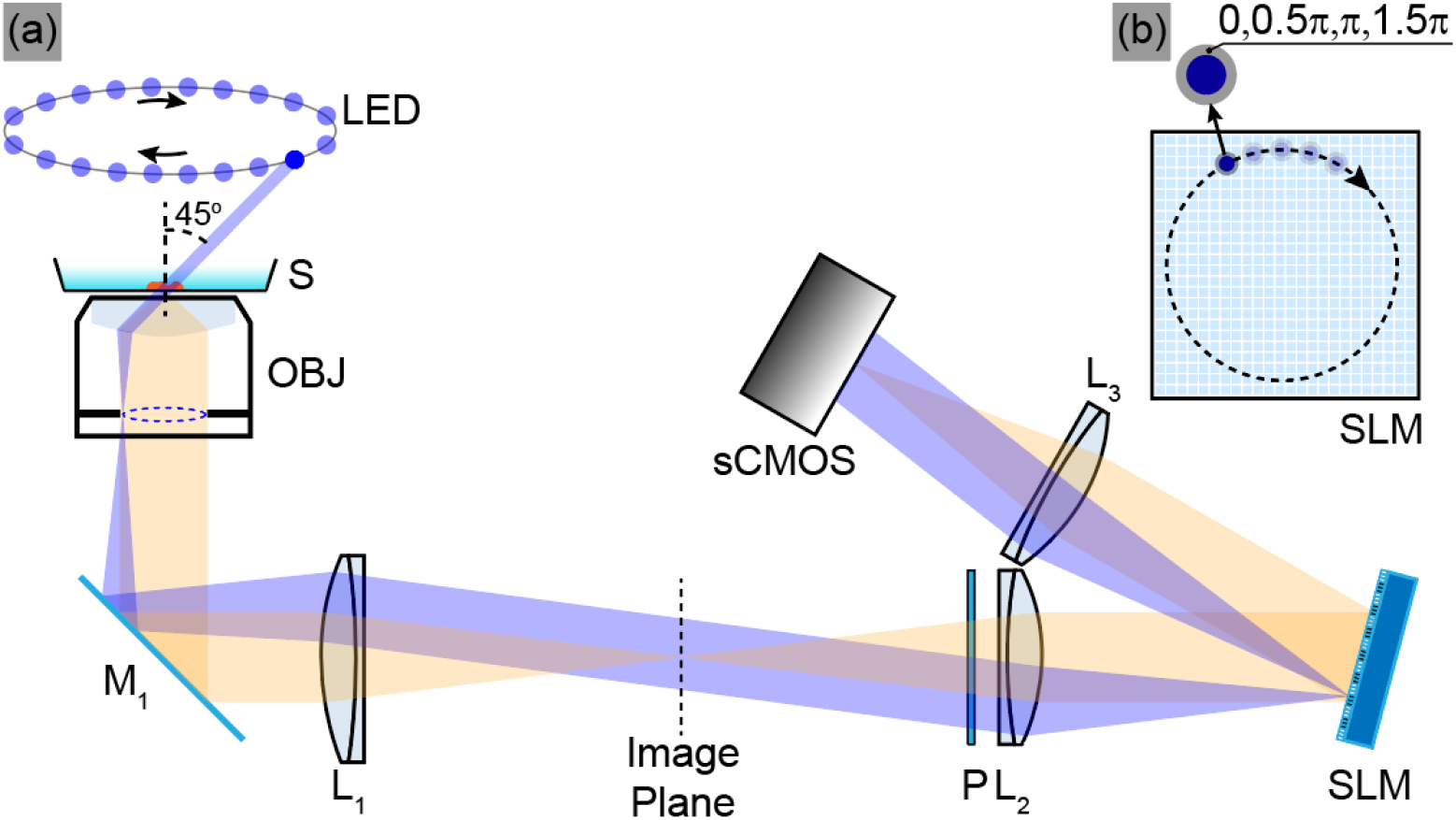
The schematic diagram of the proposed PCT. **(a)** Experimental setup of the proposed PCT with scanning illumination. **(b)** Phase modulation on the object wave spectrum by a phase-type SLM. L_1_-L_3_, achromatic lens; M, mirror; S, sample; OBJ, objective lens; P, linear polarizer; SLM, spatial light modulator.

In our PCT system, for weakly scattering samples, the object wave 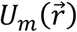under the *m*_*th*_ illumination is simply the linear summation of the 2D unscattered field 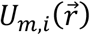 and the 2D scattering field 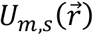 of the sample [27]. Here, 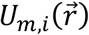 was indeed the illumination field at the focal plane of OBJ and can be expressed as 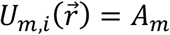 exp 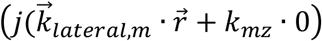 with 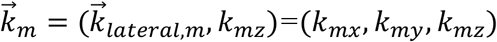being the 3D illumination vector. After the Fourier transform of the lens L_2_, the frequency spectrum of the 2D light field 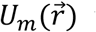 was distributed over the working plane of a phase-type SLM (MSP1920-400-800-HSP8, Meadowlark Optics, USA) located at the confocal plane of the lenses L_2_ and L_3_. The phase response calibration of the SLM can be found in Appendix 5.4. Considering the oblique illumination from an LED has an extended spectrum (focused as a spot with a diameter of 63 μm on the SLM plane), the SLM retards the phase of the unscattered term and low-frequency scattering term of the object wave (collectively called non-high-frequency term 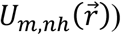with four phase values (0, 0.5π, π, and 1.5π). After being Fourier transformed by the lens L_3_, the formed phase contrast image is recorded by a sCMOS camera (Zyla 4.2, Andor, UK).

Since the phase-contrast image is the interference between the non-high-frequency term 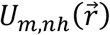 and the high-frequency scattering term 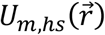, the 2D light field 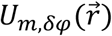reaching to the sCMOS can be expressed as follows:

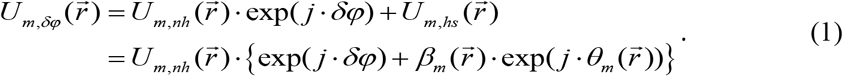

Here, 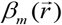 and 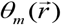 are the amplitude ratio and the phase difference between the high-frequency scattering and the non-high-frequency terms under the *m*_*th*_ illumination, respectively. *δφ* is the phase shift of 0, 0.5π, π, and 1.5π induced by the SLM on the object wave spectrum except the high-frequency scattering term.

The phase-shifted intensity images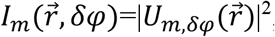, and they can be expressed as:

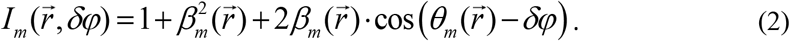

Notably, a linear polarizer P was positioned before the SLM to make the input light have a polarization direction along the fast axis of the SLM. Through simple phase-shifting processing, 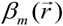 and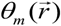 can be obtained [28]. Since the oblique illuminations from the LEDs are focused as spots with a diameter of 63 μm on the SLM plane, the sample structures with a size larger than ∼ 80 μm will be phase-modulated by the SLM. Meanwhile, in our PCT, the field of view is ∼ 100 μm × 100 μm. Consequently, the 2D high-frequency scattering term 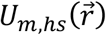 at the focal plane of OBJ can be approximately calculated as:

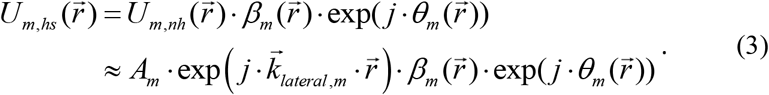

As the amplitude *A*_*m*_ and the lateral illumination vector 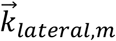 of each illumination field were determined in advance by installing a temporary lens (Appendix 5.1), the 2D high-frequency scattering term 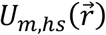 at the focal plane of OBJ was easily acquired.

According to the Fourier diffraction theorem [5], the frequency spectrum of the 2D high-frequency scattering term 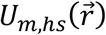 under a certain illumination angle has the following approximate relation with the 3D frequency spectrum of the sample’s scattering potential 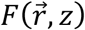:

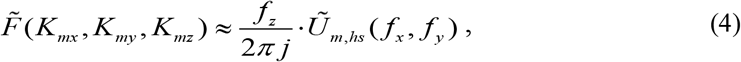

where the symbol ∼ above the functions represents the spatial Fourier transform. The 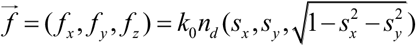 is the scattering spectrum vector with *n*_*d*_ being the RI value of the matching medium of the OBJ. As the actual system is diffraction-limited, the OBJ can only collect the scattered light within the aperture angle, *s*_*x*_ and *s*_*y*_ satisfy the relation 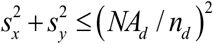 with *NA*_*d*_ being the numerical aperture of the OBJ. The 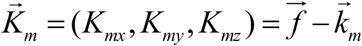 is the frequency spectrum vector of the scattering potential of the sample. By bringing Eq. (3) into Eq. (4), we finally obtained the approximate frequency spectrum of the scattering potential of the sample (spherical crown) along the *m*_*th*_ illumination:

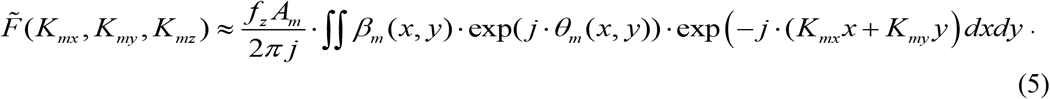

As the illumination annularly switches 25 angles, the 3D frequency spectrum distributions of the sample’s scattering potential within different Ewald crowns were determined and stitched together. Then, a Wiener filtering procedure was added to avoid the weight imbalance induced by repeatedly counting overlapped frequency components during the 3D frequency spectrum stitching. After performing the inverse Fourier transform, a 3D RI alike map of the sample can be achieved:

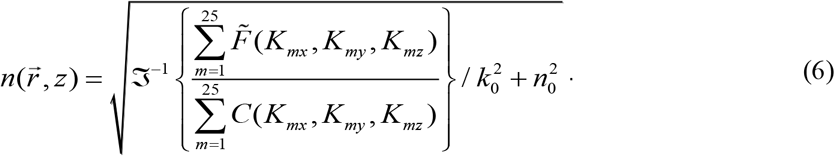

Here, 𝔍^−1^ {·} represents the inverse Fourier transform. And *C*(*K*_*mx*_, *K*_*my*_, *K*_*mz*_) is the Ewald crown under the *m*_*th*_ illumination and it equals to 1 within the effective region determined by 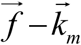 . It is worth mentioning that the SLM modulates the phase of both the unscattered term and low-frequency scattering term of the object wave, the phase of 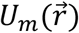 is not the correct phase of the sample but the phase difference of the high-frequency structures against the surrounding context. Once the 3D apparent-RI map of a sample is calculated with 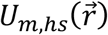 under 25 illuminations by using Eq. (5), it can still reveal the 3D distribution of high-frequency structures of a sample, despite not the 3D RI distribution of the sample. Throughout the manuscript, the obtained 3D apparent-RI distribution is normalized between 0 and 1. Compared with the ideal full aperture illumination, the frequency spectrum coverage ratio in the current PCT under 25 illuminations is 8% compared to that of 12% in the traditional ODT under 100 illuminations [9]. Therefore, reducing the number of azimuth angle at the same illumination polar angle will not lose too much frequency spectrum information, which can speed up four times under the premise of not losing too much image quality. Due to the annular oblique illumination with an effective *NA*_*illu*_ of 0.71, our PCT has a considerable spatial resolution. The theoretical lateral resolution is calculated as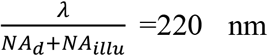, while the theoretical axial resolution is calculated as Minimum 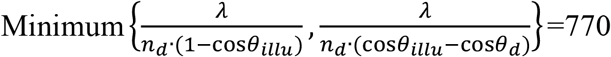 nm with θ_*illu*_ and θ_*d*_ being the llumination polar angle (45°) and the OBJ aperture angle (72°), respectively. Due to the physical dimension of the 200-nm PMs and the system aberration, the measured spatial resolution in Part 2.2 (lateral 270 ± 9 nm and axial 890 ± 42 nm) is larger than the theoretical one. In addition, the partially coherent scanning illuminations from 25 LEDs endow the system with noise-free characteristics and a higher image quality (Appendix 5.5: image quality comparison of PCT with laser-based ODT).

### 3.2. Sample preparation

#### single-layer dense 200-nm PMs

First, 10 μL stock solution containing 5% solid content was diluted into 2000 μL of deionized water in a Petri dish (FD5040-100, World Precision Instrument, USA), which was then gently shaken by hand for 2 minutes to accelerate the microsphere diffusion. The mixed solution was then incubated for 30 minutes at room temperature. The PMs were deposited and immobilized on the bottom of poly-D-lysine coated glass plate.

#### 3D sparsely distributed 200-nm PMs

The low-concentration mixed solution containing 200-nm PMs were prepared by successively adding 2000 μL of deionized water, 1 μL of a stock solution containing 5% solid content, and 200 μL of liquid agarose gel into a cone-bottom centrifugal tube. Then the mixed solution was shaken evenly by a vortex mixer at 1000 revolutions/minute for 2 minutes. Then, 10 μL of the mixed solution was dropped on a glass slide and covered by a coverslip. Before the mixed solution sandwiched between the glass and the coverslip was completely curdled, the sample was reversed up and down every 10 minutes to make the 200-nm PMs sparsely distributed.

#### Cell fixation

Before the COS7 cells were fixed with the fresh 4% formaldehyde solution, they were incubated in a 37 °C environment containing 5% CO_2_. For 1 L of 4% formaldehyde, add 800 mL of 1X PBS to a glass beaker on a stir plate in a ventilated hood. Heat while stirring to approximately 60 °C. Take care that the solution does not boil. Add 40g of paraformaldehyde powder to the heated PBS solution. The powder will not immediately dissolve into solution. Slowly raise the pH by adding 1 mol/L NaOH dropwise from a pipette until the solution clears. Once the paraformaldehyde is dissolved, the solution should be cooled and filtered. Adjust the volume of the solution to 1 L with 1X PBS. Recheck the pH and adjust it with small amounts of dilute hydrochloric acid to approximately 6.9.

## 4. Discussions and Conclusions

This work proposed the PCT by combining quantitative phase contrast microscopy (QPCM) with scanning illumination. Under each oblique illumination, a phase-type spatial light modulator (SLM) was used to modulate the zero- and low-frequency terms (non-high-frequency term) of the sample with a phase series of 0, 0.5π, π, and 1.5 π. 2D high-frequency scattering field of the sample can be recovered from the recorded phase shifted intensity images. Once 25 scattering fields are obtained sequentially after the scan of 25 illumination angles, the 2D frequency spectra of these scattering fields are mapped into the corresponding Ewald crown (the 3D frequency spectrum) of the scattering potential of the sample. Eventually, 3D structure distribution can be obtained by applying an inverse Fourier transform to the 3D frequency spectrum. The partially coherent scanning illuminations from 25 LEDs endow the system with high-resolution and noise-free characteristics. Meanwhile, the structure of common path interference makes the system very immune to external disturbances. The proposed PCT was successively applied to render the 3D structure distributions of single-layer dense 200-nm PMs, 3D sparsely distributed 200-nm PMs, and LDs in a fixed COS7 cell to verify its feasibility and performance. These confirmatory experiments showed that the PCT can provide a high-quality 3D structure distribution for a transparent sample. Subsequently, the proposed PCT was applied to visualize the 3D structures of sub-organelles inside COS7 cells. Therefore, the PCT has more comprehensive advantages and broader application prospects in many fields.

However, the actual phase modulation not only occurs to the unscattered field but also the low-frequency components of a sample due to the fact that the LEDs have an extended spectrum and the phase-modulated area is larger than the real unscattered component of a sample. Therefore, the recovered 2D scattering field in our method is indeed a high-frequency one, meaning that the 3D RI distribution obtained by our method is inconsistent with the true value. As a result, our method can reveal the 3D structure distribution of a sample by mapping a series of 2D high-frequency scattering fields, however, not the 3D RI distribution. In the future, fiber-based partially coherent illumination can be used to decrease the size of the focused illumination fields so that only the unscattered field is phase-modulated.

Notably, for strong scattering samples, like biological tissue, the generated unscattered field energy is very small and even barely exists, thus the corresponding phase modulation is not valid. Therefore, the PCT is not suitable for thick strong scattering samples. Theoretically, the more scanning angles, the better the 3D imaging. However, the large number of LED switches is unconducive to the imaging speed. Consequently, we compromise to set the number of LEDs to 25 in the experiments. In the future, the imaging speed and the full aperture illumination of the proposed PCT will be ensured by digital scanning through a DMD or an SLM.

## 5. Appendix

### 5.1 Illumination vector calibration

Determining the 2D high-frequency scattering field 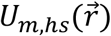 in Eq. (3) needs to know the amplitude *A*_*m*_ and the lateral illumination vector 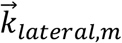 of the illumination field. For this purpose, we developed an effective and robust approach to determine the illumination field. First, a temporary lens was installed between the sCMOS and the SLM to image the working plane of the SLM and the rear focal plane of the OBJ simultaneously. Second, a strong scattering sample, like a cured photoresist, was installed at the focal plane of the OBJ, and the sCMOS synchronously recorded the intensity images when the LEDs were turned on one by one. Therefore, the pupil aperture of the OBJ and its center were accurately determined after adding these intensity images (**Fig. 7(a)**). Third, the sample to be investigated was installed at the focal plane of the OBJ, and the sCMOS also synchronously recorded the phase-shifted intensity images when the LEDs were turned on sequentially. **Figure 7(b)** illustrates the image of the pupil plane of the OBJ under a certain illumination angle, demonstrating that the illumination field was focused as a spot. Therefore, the lateral illumination vector 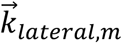 was accurately determined as (*x*_*m*_-*x*_0_, *y*_*m*_-*y*_0_)·*k*_0_·*NA*_*d*_/*r* with *NA*_*d*_ being the numerical aperture of the OBJ and *r* being the radius of the pupil aperture of the OBJ. Moreover, the illumination vector 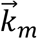 was thereby obtained as 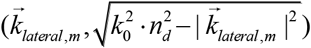 with *n*_*d*_ being the RI value of the matching medium of the OBJ. The amplitude *A*_*m*_ was determined by normalizing the intensity values of the focused spots at the pupil plane of the OBJ under the different illumination angles. In addition, through this temporary lens, the phase patterns loaded onto the SLM for phase-modulating the unscattered fields were well-matched with these focused illumination fields. Consequently, the 2D high-frequency scattering field 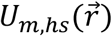 at the focal plane of OBJ was easily acquired.

**Fig. 7.**
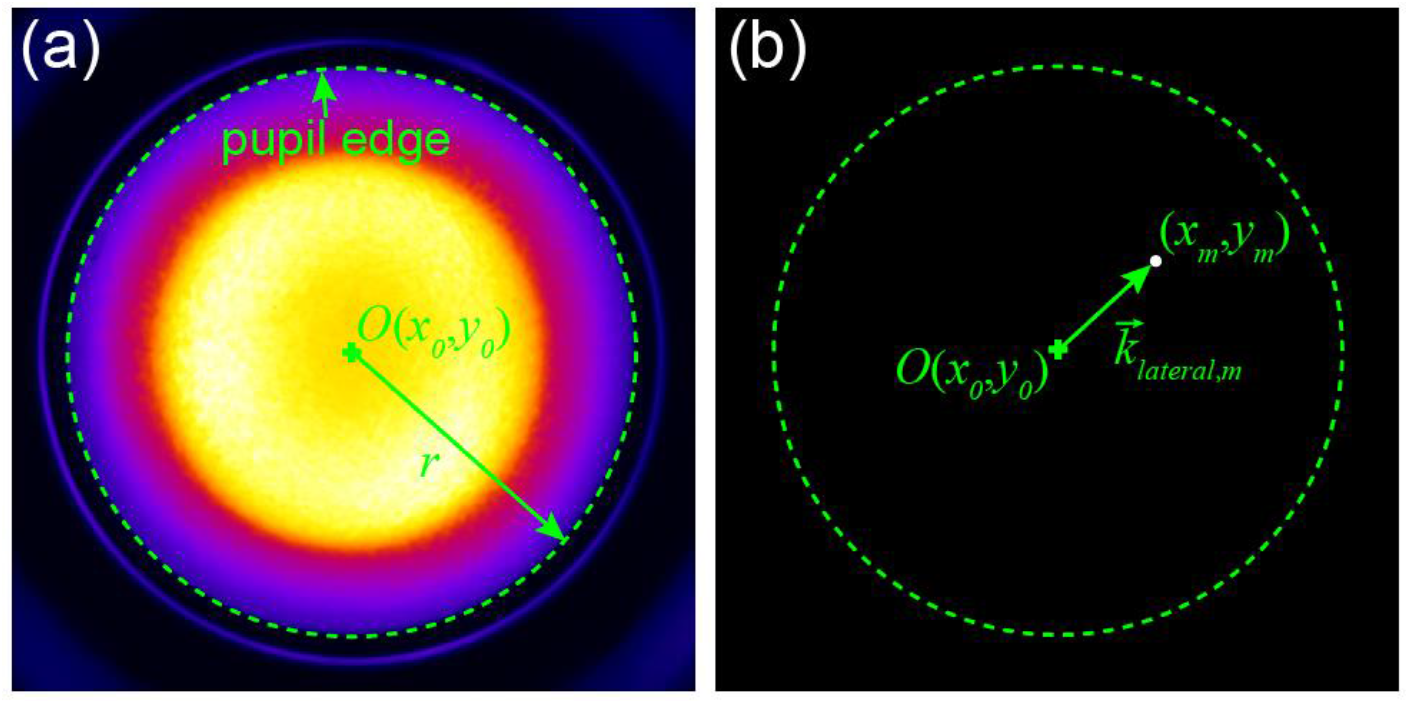
Illumination vector calibration. **(a)** Determination of the pupil aperture of the objective lens and its center. **(b)** Determination of the amplitude of the illumination field and the lateral illumination vector. *r* is the radius of the pupil aperture of the OBJ.

### 5.2. Illumination stability

As described in the Methods Part, at the beginning of the data acquisition in PCT, the illumination vectors are determined by installing a temporary lens between the sCMOS and the SLM. The stability of the illumination vectors is crucial for the high-quality 3D reconstruction of PCT. Therefore, we tested the repeatability of illumination vectors against the external disturbs and the vibration of rotating module. We collected and calibrated 25 angles of illumination vectors at 10-minute intervals over 100 minutes. The test results are shown in **Fig. 8**, where the blue dots correspond to illumination vectors at different time points. And the inset in **Fig. 8** further shows the variation of the illumination vector at a certain angle, revealing that the variation is very tiny in the proposed PCT. Compared with the reported ODT technique [9], the variation of the illumination vector in PCT is reduced by an order of magnitude.

**Fig. 8.**
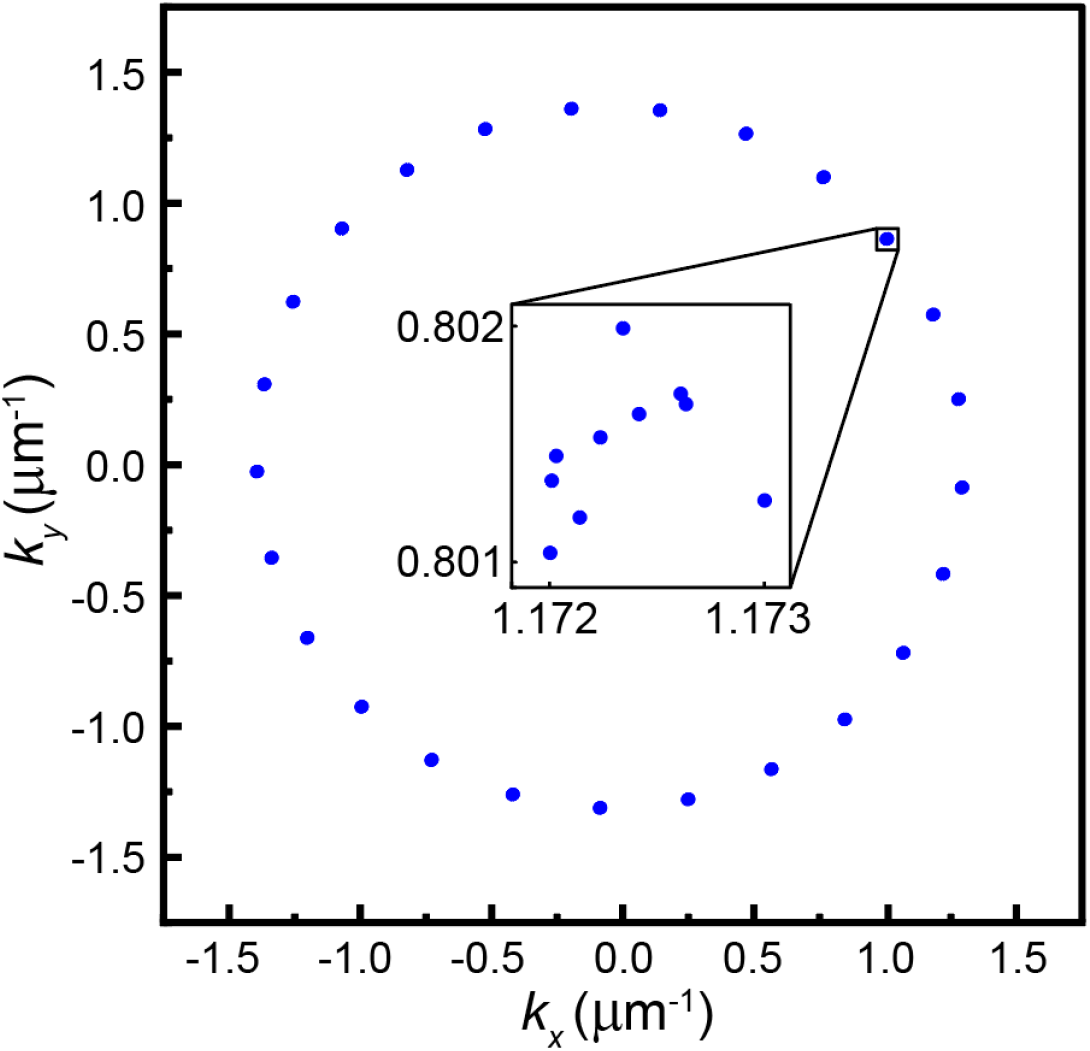
Deviation of illumination vectors at different time points.

### 5.3. Parallel control algorithm

A high-speed data acquisition card (USB-6363, National Instruments, USA) is used to synchronously control 25 individual LEDs, the SLM, and the sCMOS camera. The control sequence diagram of the PCT is shown in **Fig. 9**. 25 LEDs are turned on and off in sequence, during which, the SLM and the sCMOS camera are triggered four times (four phase modulations) when each LED is turned on. For the SLM, to avoid the pattern loading time, 100 phase modulation patterns (four phase modulations for each of 25 LEDs) have been loaded into the SLM cache in advance, and the displayed pattern on the SLM’s working plane for retarding the phase of the unscattered term and low-frequency term of the object wave under a certain illumination angle is switched by a rising edge signal. The switch time of the displayed pattern of the SLM is on the order of 1 μs, which can be negligible. The sCMOS camera immediately records the intensity distribution before the next switch of the displayed pattern of the SLM. The recording of the sCMOS camera includes two parts, the global exposure of the selected pixels and the signal readout, as shown in **Fig. 9**. The global exposure is determined by the interval between a rising and a falling edge signal, which is limited by the illumination intensity of the LEDs. In our PCT, the global exposure time can reach 1 *ms*. The signal readout is triggered by a falling edge signal, and its duration is determined by the row number of the selected pixel area due to the rolling shutter mode of the Andor camera (∼ 10 *ms* for 1080 rows). Therefore, the time it takes for the sCMOS camera to record once is expressed as 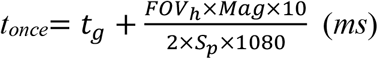 (*ms*) with *t*_*g*_, *FOV*_*h*_, *Mag*, and *S*_*p*_ being the global exposure time, the height of the imaging field of view of the sample, the system magnification, and the pixel size of the sCMOS camera, respectively. Notably, every rising edge signal for the SLM is synchronized with a corresponding rising edge signal for the sCMOS camera. The duration of each LED is four times the interval between adjacent two rising edge signals for the SLM or the sCMOS camera due to four phase modulations. With 100 phase modulation patterns being successively displayed on the working plane of the SLM according to the illumination angle and the phase modulation, the sCMOS camera synchronously records 100 intensity distributions that are used for reconstructing the 3D quasi-RI distribution of the sample. Therefore, the frame rate of the PCT can be normally expressed as 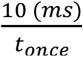 (fps).

**Fig. 9.**
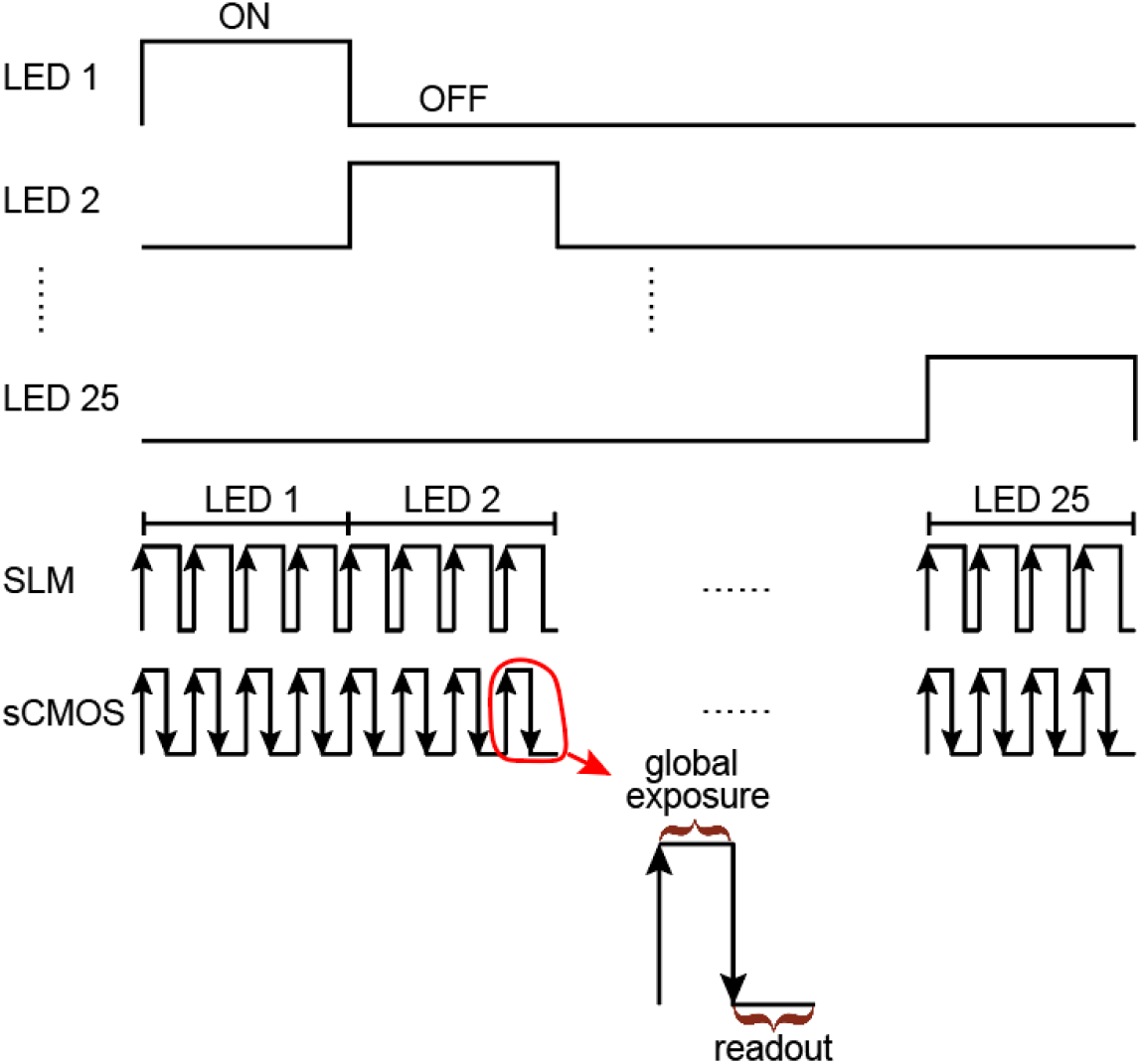
Control sequence diagram of the PCT.

### 5.4. SLM calibration

SLM is the key component for recovering the 2D scattering field of the sample in PCT, and it needs a calibration between the generated phase and the applied voltage since there exists a nonlinear optical response of liquid crystal to loaded voltage. A simple method is used to accurately calibrate the SLM, which measures the response property of each pixel and avoids the wavefront curvature induced by optical elements [29]. As is known, the liquid crystals in SLMs only modulate the light with the polarization direction along the fast axis of the SLM (termed as y-axis), while it only acts as a mirror to that along the slow axis (termed as x-axis). Based on this, we provisionally add an additional linear polarizer between the SLM and the sCMOS and set its polarization direction 45° with respect to the *x*-axis. Meanwhile, the original linear polarizer P in **Fig. 6(a)** is rotated to make its polarization direction 45° with respect to the *x*-axis too. Then a used LED is positioned before the original polarizer P and emits divergent light beam, which traverses through the original polarizer P and is modulated by the SLM. Therefore, the resulted light beams along the *x* and *y* axes are written as

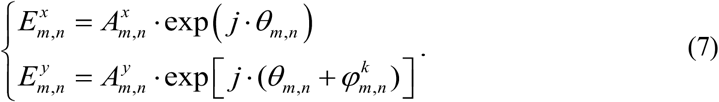

Here, *m* and *n* represent the pixel coordinates of the SLM along the *x* and *y* directions.

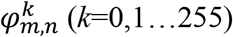 is the generated phase value at a certain pixel when the *k*_*th*_ voltage is added. Then, the light intensity induced at a certain pixel after passing through the additional linear polarizer is calculated as:

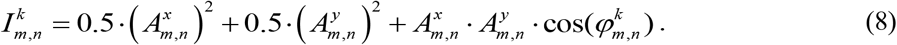

The dc term 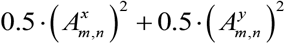 and the modulation depth 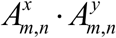 can be determined when applying the voltage from 0 to the maximal voltage allowed and determining the maximal and minimal intensities for the intensity series obtained. Then the relation between the induced phase value and the added voltage is obtained with the cos^-1^{·} function, as is shown with the red curve in **Fig. 10**. Notably, though 25 LEDs have the same model, there is a slight difference in the SLM response curve under their illuminations. Therefore, the SLM response calibration is carried out on 25 LEDs, respectively.

**Fig. 10.**
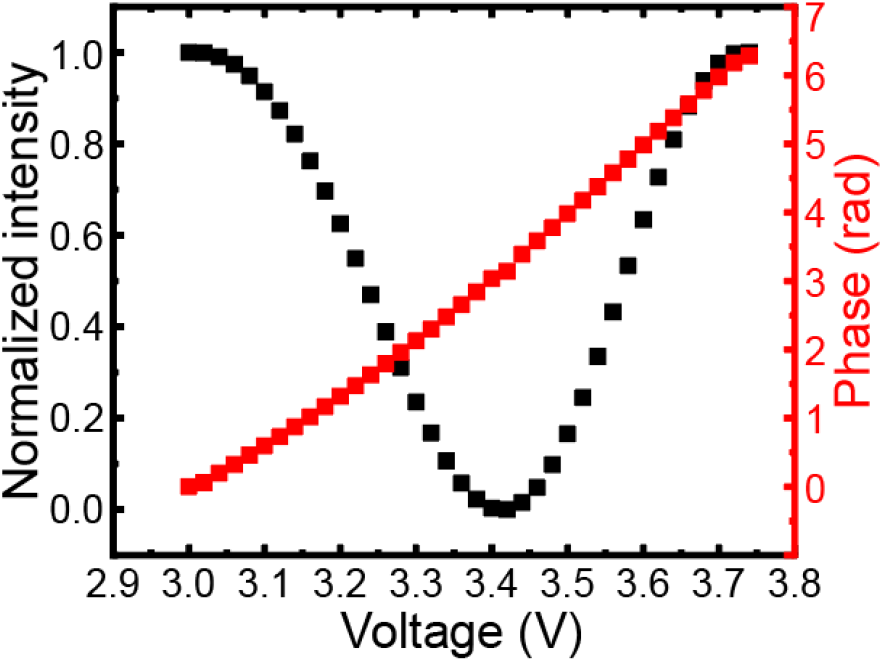
SLM response calibration for a certain LED: the measured intensity (black) and the modulated phase (red) versus the loaded voltage.

### 5.5. Image quality comparison

As mentioned before, the partially coherent illuminations from 25 LEDs in the proposed PCT endow the system with excellent image quality. And to intuitively reflect this advantage, we compared the image quality of empty areas reconstructed by the proposed PCT, the laser scanning-based PCT, and the structured illumination-based ODT (SI-ODT) [30], respectively. In the laser scanning-based PCT, only the illumination part is changed by a laser coupled into a rotating module (RM) compared to the proposed PCT. The schematic diagram of the laser scanning illumination is shown in **Fig. 11(a)**. The emitted light from a laser was transmitted by a single mode fiber (SMF) that was positioned at the focal point of the lens L_4_, so that the emitted coherent light was collimated into a plane wave. After being spatially confined by a diaphragm D, the collimated light was scanned by the RM containing mirrors M_2_ and M_3_. The RM was installed 5 cm above the sample positioned at the focal plane of the objective lens (OBJ; 100×/1.44, Leica, Germany. As the RM rotated, the sample was sequentially illuminated in 25 directions, which share a polar angle of 45° and differentiate with an azimuth angle of 2mπ/25 with *m*=0, 2, …, 24. The phase modulations on the object wave spectrum under the laser illuminations and the 3D RI-alike reconstruction of the sample are the same as the proposed PCT. The SI-ODT system uses a coherent laser and a DMD to generate the structured illumination, and an off-axis reference beam is used to record the holograms. The details of the SI-ODT system can be found in [30]. **Figures 11(b)-(d)** are the normalized RI histograms of empty areas reconstructed by PCT, laser scanning-based PCT (Laser-PCT), and SI-ODT, respectively. The normalized RI histograms reflect the background uniformity. And the narrower the distribution of the histograms, the higher the image quality. From **Figs. 11(b)-(d)** we can see that the partially coherent illuminations from 25 LEDs provide a better image quality compared to the laser-based illumination. And the image quality of SI-ODT is a little better than that of Laser-PCT due to the multiple harmonic illumination induced by the DMD.

**Fig. 11.**
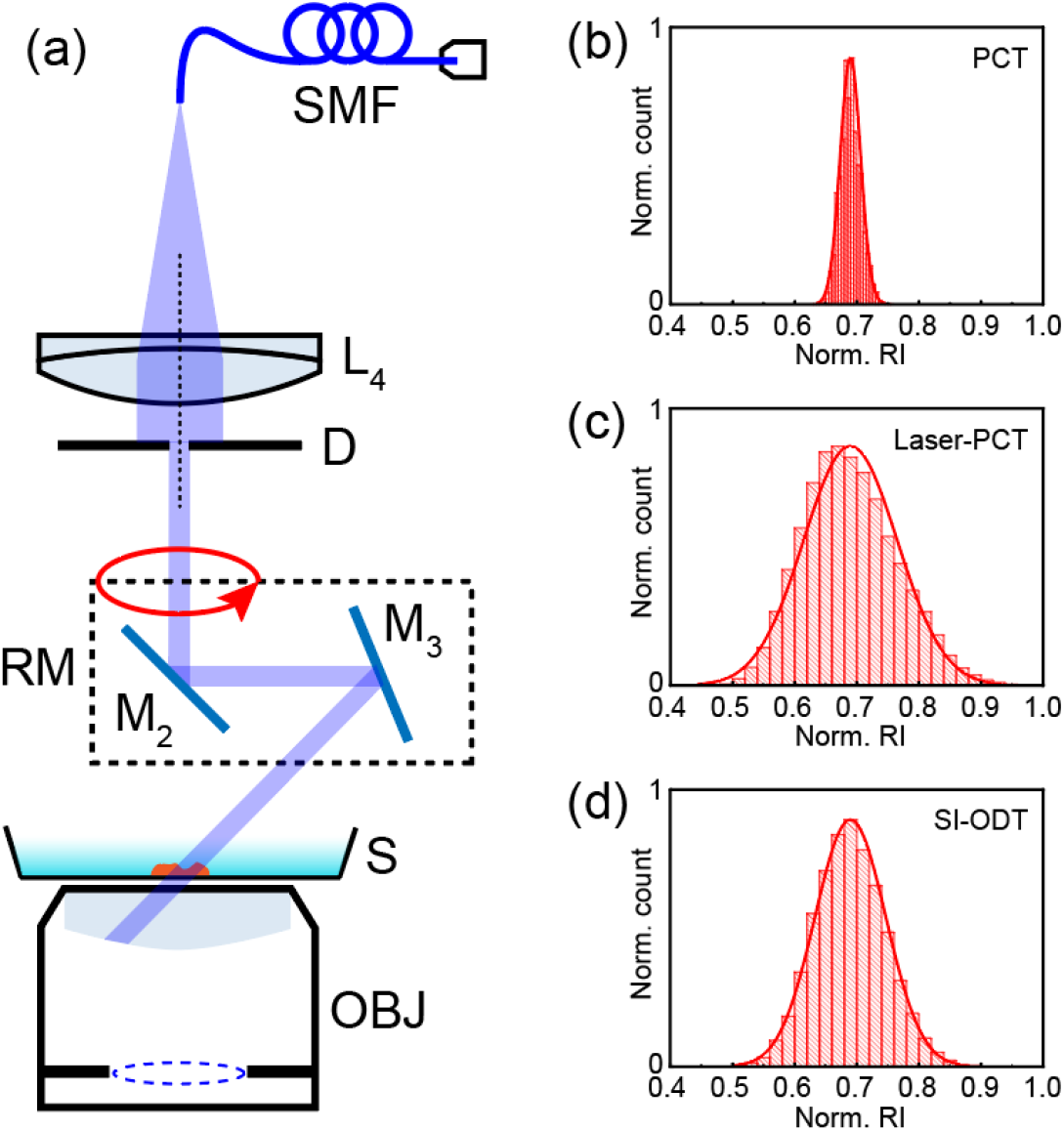
Image quality comparison. (a) Schematic diagram of laser scanning illumination. (b)-(d) Normalized RI histograms of empty areas reconstructed by PCT, laser scanning-based PCT, and SI-ODT, respectively. D, diaphragm; L_4_, achromatic lens; M, mirror; RM, rotating module containing two mirrors M_2_ and M_3_; S, sample; SMF, single mode fiber; OBJ, objective lens.

## Acknowledgments

Y. M. and W. J. F. performed experiments and data analysis. Y.

Z. L. completed the control of the system. L. M., J. J. Z., S. A., and M. L. contributed to data analysis. Y. M. wrote the draft of the manuscript. P. G. supported the subject and revised the manuscript. All the authors edited the manuscript.

## Data availability

Data underlying the results presented in this paper are not publicly available at this time but may be obtained from the authors upon reasonable request.

## References

1. F. Zernike, “Phase contrast, a new method for the microscopic observation of transparent objects,” Physica. 9, 686–698 (1942).

2. C. S. Anderson, “Fringe visibility, irradiance, and accuracy in common path interferometers for visualization of phase disturbances,” Appl. Opt. 34(32), 7474–7485 (1995).

3. H. Kadono, M. Ogusu, and S. Toyooka, “Phase shifting common path interferometer using a liquid-crystal phase modulator,” Opt. Commun. 110, 391–400 (1994).

4. X. Chen, M. E. Kandel, and G. Popescu, “Spatial light interference microscopy: principle and applications to biomedicine,” Adv. Opt. Photonics. 13(2), 353–425 (2021).

5. Y. J. Sung, W. Choi, C. F. Yen, K. Badizadegan, R. R. Dasari, and M. S. Feld, “Optical diffraction tomography for high resolution live cell imaging,” Opt. Express. 17(1), 266–277 (2009).

6. K. Kim, H. O. Yoon, M. D. Silva, M. Dao, R. R. Dasari, and Y. K. Park, “High-resolution 3D imaging of red blood cells parasitized by Plasmodium falciparum and in situ hemozoin crystals using optical diffraction tomography,” J. Biomed. Opt. 19(1), 011005 (2014).

7. T. K. Kim, B. W. Lee, F. Fujii, K. H. Lee, S. Lee, Y. K. Park, J. K. Kim, S. W. Lee, and C. G. Pack, “Mitotic chromosomes in live cells characterized using high-speed and label-free optical diffraction tomography,” Cells. 8(11), 1368 (2019).

8. Y. Cotte, F. Toy, P. Jourdain, N. Pavillon, D. Boss, P. Magistretti, P. Marquet, and C. Depeursinge, “Marker-free phase nanoscopy,” Nat. Photonics. 7,113–117 (2013).

9. D. S. Dong, X. S. Huang, L. J. Li, H. Mao, Y. Q. Mo, G. Y. Zhang, Z. Zhang, J. Y. Shen, W. Liu, Z. M. Wu, G. H. Liu, Y. M. Liu, H. Yang, Q. H. Gong, K. B. Shi, and L. Y. Chen, “Super-resolution fluorescence-assisted diffraction computational tomography reveals the 3D landscape of the cellular organelle interactome,” Light-Sci. Appl. 9, 11 (2020).

10. K. R. Lee, K. Kim, G. Kim, S. Shin, and Y. K. Park, “Time-multiplexed structured illumination using a DMD for optical diffraction tomography,” Opt. Lett. 42(5), 999–1002 (2017).

11. J. Oh, J. S. Ryu, M. Lee, J. Jung, S. Y. Han, H. J. Chung, and Y. Park, “3D label-free observation of individual bacteria upon antibiotic treatment using optical diffraction tomography,” Biomed. Opt. Express. 11(3), 1257–1267 (2020).

12. M. Lee, K. Kim, J. Oh, and Y. K. Park, “Isotropically resolved label-free tomographic imaging based on tomographic moulds for optical trapping,” Light-Sci. Appl. 10(1), 102 (2021).

13. K. R. Lee, S. Shin, Z. Yaqoob, P. T. C. So, and Y. K. Park, “Low-coherent optical diffraction tomography by angle-scanning illumination,” J. Biophotonics. 12(5), e201800289 (2019).

14. C. Park, K. R. Lee, Y. Baek, and Y. K. Park, “Low-coherence optical diffraction tomography using a ferroelectric liquid crystal spatial light modulator,” Opt. Express. 28(26), 39649–39659 (2020).

15. S. Chowdhury, W. J. Eldridge, A. Wax, and J. A. Izatt, “Refractive index tomography with structured illumination,” Optica. 4(5), 537–537 (2017).

16. B. Bhaduri, H. Pham, M. Mir, and G. Popescu, “Diffraction phase microscopy with white light,” Opt. Lett. 37(6), 1094–1096 (2012).

17. M. L. Zhang, Y. Ma, Y. Wang, K. Wen, J. J. Zheng, L. X. Liu, and P. Gao, “Polarization grating based on diffraction phase microscopy for quantitative phase imaging of paramecia,” Opt. Express. 28, 29775–29787 (2020).

18. T. H. Nguyen, M. E. Kandel, M. Rubessa, M. B. Wheeler, and G. Popescu, “Gradient light interference microscopy for 3D imaging of unlabeled specimens,” Nat. Commun. 8, 210 (2017).

19. G. Vishnyakov, G. Levin, V. Minaev, M. Latushko, N. Nekrasov, and V. Pickalov, “Differential interference contrast tomography,” Opt. Lett. 41(13), 3037–3040 (2016).

20. P. Zdańkowski, J. Winnik, K. Patorski, P. Gocłowski, M. Ziemczonok, M. Józwik, M. Kujawińska, and Maciej Trusiak, “Common-path intrinsically achromatic optical diffraction tomography,” Biomed. Opt. Express. 12(7), 4219–4234 (2021).

21. J. M. Soto, J. A. Rodrigo, and T. Alieva, “Partially coherent illumination engineering for enhanced refractive index tomography,” Opt. Lett. 43(19), 4699–4702 (2018).

22. J. J. Li, A. C. Matlock, Y. Z. Li, Q. Chen, C. Zuo, and L. Tian, “High-speed in vitro intensity diffraction tomography,” Advanced Photonics. 1(6), 066004 (2019).

23. C. Michael, R. David, L. H. Yuan, C. Shwetadwip, and W. Laura, “Multi-layer Born multiple-scattering model for 3D phase microscopy,” Optica. 7, 394–403 (2020).

24. J. Zhao, A. Matlock, H. B. Zhu, Z. Q. Song, J. B. Zhu, B. Wang, F. K. Chen, Y. W. Zhan, Z. C. Chen, Y. H. Xu, X. C. Lin, L. Tian, and J. X. Cheng, “Bond-selective intensity diffraction tomography,” Nat. Commun. 13(1), 7767 (2022).

25. J. J. Li, N. Zhou, J. S. Sun, S. Zhou, Z. D. Bai, L. P. Lu, Q. Chen, and C. Zuo, “Transport of intensity diffraction tomography with non-interferometric synthetic aperture for 3D label-free microscopy,” Light-Sci. Appl. 11(1), 154 (2022).

26. A. B. Ayoub, A. Roy, and D. Psaltis, “Optical diffraction tomography using nearly in-line holography with a broadband LED source,” Appl. Sci. 12(3), 951 (2022).

27. T. Kim, R. J. Zhou, M. Mir, S. D. Babacan, P. S. Carney, L. L. Goddard, and G. Popescu, “White-light diffraction tomography of unlabelled live cells,” Nat. Photonics. 8(3), 256–263 (2014).

28. Y. Ma, T. Q. Dai, Y. Z. Lei, J. J. Zheng, M. Liu, B. D. Sui, Z. J. Smith, K. Q. Chu, L. Kong, and P. Gao, “Label-free imaging of intracellular organelle dynamics using flat-fielding quantitative phase contrast microscopy (FF-QPCM),” Opt. Express. 30(6), 9505–9520 (2022).

29. Y. Ma, L. Ma, J. J. Zheng, M. Liu, Z. Zalevsky, and P. Gao, “High spatio-temporal resolution condenser-free quantitative phase contrast microscopy,” Front. Phys.-Lausanne. 10, 892529 (2022).

30. R. Liu, K. Wen, J. Li, Y. Ma, J. Zheng, S. An, J. Min, Z. Zalevsky, B. Yao, and P. Gao, “Multi-harmonic structured illumination-based optical diffraction tomography,” Appl. Opt. 62(35), 9199–9206 (2023)..

